# Use of Stem Cell-Derived Cardiomyocyte and Nasal Epithelium Models to Establish a Multi-Tissue Model Platform to Validate Repurposed Drugs Against SARS-CoV-2 Infection

**DOI:** 10.1101/2024.05.22.595397

**Authors:** Nathan J Gödde, Carmel M O’Brien, Elizabeth Vincan, Aditya Vashi, Stephanie Olliff, Bang M Tran, Shafagh A Waters, Sarah Goldie, Petrus Jansen van Vuren, Shane Riddell, Matthew P Bruce, Vinti Agarwal, Eugene Athan, Kim R Blasdell, Simran Chahal, Darren J Creek, Faheem, Hardik A Jain, Carl M Kirkpatrick, Anupama Kumar, Christopher A MacRaild, Mohammed Muzaffar-Ur-Rehman, Murugesan Sankaranarayanan, Rohan M Shah, Ian K Styles, Mary Tachedjian, Natalie L Trevaskis, Nagendrakumar B Singanallur, Alexander J McAuley, Seshadri S Vasan

## Abstract

The novel coronavirus disease (COVID-19) and any future coronavirus outbreaks will require more affordable, effective and safe treatment options to complement current ones such as *Paxlovid*. Drug repurposing can be a promising approach if we are able to find a rapid, robust and reliable way to down-select and screen candidates using *in silico* and *in vitro* approaches. With repurposed drugs, ex vivo models could offer a rigorous route to human clinical trials with less time invested into nonclinical animal (*in vivo*) studies. We have previously shown the value of commercially available *ex vivo/3D* airway and alveolar tissue models, and this paper takes this further by developing and validating human nasal epithelial model and embryonic stem cells derived cardiomyocyte model. Five shortlisted candidates (fluvoxamine, everolimus, pyrimethamine, aprepitant and sirolimus) were successfully compared with three control drugs (remdesivir, molnupiravir, nirmatrelvir) when tested against key variants of the SARS-CoV-2 virus including Delta and Omicron, and we were able to reconfirm our earlier finding that fluvoxamine can induce antiviral efficacy in combination with other drugs. Scalability of this high-throughput screening approach has been demonstrated using a liquid handling robotic platform for future ‘Disease-X’ outbreaks.

## Introduction

Almost five years after the start of the COVID-19 pandemic, there remain few effective antiviral drugs for its treatment. Most drugs that are available (e.g. Paxlovid) are prohibitively expensive for low-income countries, leaving them with no therapeutic interventions to treat patients. The repurposing of existing drugs can be a useful approach for the identification of deployable therapies but, given that there are several thousands of prescription drug products to choose from, selecting for antiviral efficacy using an appropriate model(s) is challenging [1]. Such efforts are not only useful for the treatment of COVID-19, but also prepare for future coronavirus outbreaks.

Early in the pandemic, high-throughput drug screening was performed using permissive cell lines, such as Vero E6, which identified a number of drug candidates with possible antiviral efficacy against SARS-CoV-2 infection (measured through the inhibition of observable cytopathic effect) [2-6]. However, many, if not most, of these drugs failed to show efficacy in more complex testing settings. Immortalised, continuous cell lines are either intrinsically tumorigenic or have been engineered as such, resulting in aberrant cellular process such as dysregulated proteosomes, impaired immune responses, and a loss of cellular polarity [7, 8]. As such, these cells behave significantly differently to those in host tissues, and frequently result in false positive or false negative results. Indeed, the lack of clinical efficacy for both chloroquine/hydroxychloroquine and ivermectin, despite strong in vitro evidence, serve as cautionary tales for the over-interpretation of antiviral efficacy based on tissue culture studies [4, 5, 9, 10].

3D tissue models, whether air-liquid interface (ALI) cultures or organoids, constitute more advanced *in vitro/ex vivo* systems for drug assessment as they overcome many of the aforementioned limitations. 3D tissue models have the advantage that they are derived from non-immortalised cells (human or animal) and are differentiated to generate cellular structures that better recapitulate human and animal tissues than is possible with immortalised cells [7], however they still have some limitations including the time and cost associated with differentiation, the need to source new seed material, and significant batch-to-batch variation. Nevertheless, their response to infection and treatment better reflects that seen in patients and, as such, may generate more reliable data for known and emerging infectious diseases.

We have recently employed an *in silico* approach to down-select nine approved drugs (plus three anti-SARS-CoV-2 control drugs) for assessment using a commercially-sourced ALI model of the human airway epithelium [11-13]. Five drugs that showed promise were further evaluated against both Delta and Omicron variants of concern (VOC), with fluvoxamine exhibiting anti-SARS-CoV-2 activity, albeit at concentrations higher than can be achieved in patients [13].

SARS-CoV-2 does not, however, only affect the airway. It can infect and damage cardiac tissues, cause intestinal dysregulation, and affect cognitive functions [14-16]. Infection of nasal epithelial cells can result in prolonged viral shedding, which likely contributes to human-to-human transmission [17, 18]. Accordingly, potential drugs should show efficacy in multiple relevant tissue models before conclusions are drawn about whether to advance to *in vitro* or clinical studies. This means that a panel of models should be established that appropriately reflect the clinical tropism of a given virus.

This study builds upon our previous human airway model-based assessment of approved drugs by characterising SARS-CoV-2 infection in human nasal epithelial (HNE) and cardiomyocyte models, as well as re-assessing the previously-characterised approved drugs against both Delta and Omicron VOC in the cardiomyocytes. Furthermore, we demonstrate that the differentiated cardiomyocytes can be processed using liquid handling robotic platforms to permit their usage for high-throughput screening of antiviral compounds during outbreak conditions.

## Materials and Methods

### Human Ethics Approvals

For the generation of human nasal epithelial cells, study approval was received from the Sydney Children’s Hospital Network Ethics Review Board (HREC/16/SCHN/120) and the Medicine and Dentistry Human Ethics Sub-Committee, University of Melbourne (HREC/2057111). Written consent was obtained from all participants prior to collection of biospecimens.

Usage of human embryonic stem cells (for cardiomyocyte production), and infection of both sets of tissue models was covered by approval from the CSIRO Health and Medical Human Research Ethics Committee (2021_83_LR).

### Development of Human Nasal Epithelial Model (HNE)

De-identified, cryopreserved human nasal epithelial cells were received from the Molecular and Integrative Cystic Fibrosis Research Centre (University of New South Wales, Sydney, NSW, Australia), where they were harvested from nasal turbinate brush samples followed by culture under conditional reprogram conditions as previously described [19, 20].

Mucociliary differentiation at the air-liquid interface was performed as previously described [20]. Briefly, cryovials of cells were thawed and seeded onto 6.5mm transwell inserts (Corning, Kennebunk, ME, USA) pre-coated with collagen Type I (PureCol-S; Advanced BioMatrix, San Diego, CA, USA). Cells were incubated for 7 days submerged in PneumaCult-ExPlus (STEMCELL Technologies, Vancouver, BC, Canada). After the 7-day culture, the PneumaCult-ExPlus media was substituted for PneumaCult-ALI medium (STEMCELL Technologies), and ALI differentiation was initiated by exposing the apical surface to air. Medium in the basal compartment was replaced three times per week for four weeks until differentiation was complete.

### Development of Cardiomyocyte Model

WA09 embryonic stem cells (WiCell, Madison, WI, USA) were expanded and maintained at 37°C/5% CO_2_ in mTeSR-E8 culture medium (STEMCELL Technologies) on Nunclon cultureware coated with LDEV-free Geltrex (Thermo Fisher Scientific, Waltham, MA, USA). At approximately 80% confluence, cells were harvested using ReLeSR dissociation reagent (STEMCELL Technologies) and passaged for a minimum of three times before commencing differentiation into cardiomyocytes.

For generation of cardiomyocytes, dissociated WA09 cells were plated onto Matrigel-coated 75cm^2^ tissue culture flasks (Corning) in medium supplemented with 10 μM Rho kinase inhibitor (Y-27632; STEMCELL Technologies) at 1.35×10^6^ cells/cm^2^ to achieve 95% confluence after 48h culture (Differentiation Day 0). The culture medium was replaced 24h post-seeding, and then sequentially with STEMdiff Cardiomyocyte Differentiation kit components, per the manufacturer’s protocol (STEMCELL Technologies). Following 15 days of differentiation, approximately 90% of the cells were observed to be rhythmically beating. The cultures were maintained in STEMdiff Cardiomyocyte Maintenance Medium (STEMCELL Technologies) for a further 5-7 days. Prior to infection, the cardiomyocyte cultures were dissociated using a cardiomyocyte dissociation kit (STEMCELL Technologies) before being replated in matrigel-coated 12 well plates at a seeding density of 0.5×10^6^ cells/well, resulting in a consistent surface coverage of cardiomyocytes within and between wells. Infections occurred between Day 20 and 25 of differentiation.

### Virus Stocks and Viral Titration

Virus stocks used in our previous study, [13], were used again for this work. In brief, Delta (B.1.617.2; hCoV-19/Australia/VIC18440/2021; EPI_ISL_1913206) and Omicron BA.1.1 (hCoV-19/Australia/VIC28585/2021; EPI_ISL_7771171) SARS-CoV-2 VOC were kindly provided by Drs Caly and Druce at the Victorian Infectious Diseases Reference Laboratory. Working stocks of each were grown in Vero E6 cells (American Type Culture Collection, Manassas, VA, USA), with Dulbecco’s Minimum Essential Medium (DMEM) supplemented with 2% FBS, 2 mM GlutaMAX supplement, 100 U/mL penicillin, and 100 μg/mL streptomycin (all components from Thermo Fisher Scientific). Diluted inoculum was used to inoculate Vero E6 cells for 1 h at 37 °C/5% CO_2_ before additional media was added to the flask. The flasks were incubated for 48 h before supernatant was centrifuged at 2000× g for 10 min to clarify, harvested and stored in 1 mL aliquots at −80 °C.

Identity of virus stocks was confirmed by next-generation sequencing using a MiniSeq platform (Illumina, Inc.; San Diego, CA, USA). In brief, 100 µL cell culture supernatant from infected Vero E6 cells was combined with 300 µL TRIzol reagent (Thermo Fisher Scientific) and RNA was purified using a Direct-zol RNA Miniprep kit (Zymo Research, Irvine, CA, USA). Purified RNA was further concentrated using an RNA Clean-and-Concentrator kit (Zymo Research), followed by quantification on a DeNovix DS-11 FX Fluorometer. RNA was converted to double-stranded cDNA, ligated then isothermally amplified using a QIAseq FX single cell RNA library kit (Qiagen, Hilden, Germany). Fragmentation and dual-index library preparation was conducted with an Illumina DNA Prep, Tagmentation Library Preparation kit. Average library size was determined using a Bioanalyser (Agilent Technologies, San Diego, CA, USA) and quantified with a Qubit 3.0 Fluorometer (Thermo Fisher Scientific). Denatured libraries were sequenced on an Illumina MiniSeq using a 300-cycle Mid-Output Reagent kit as per the manufacturer’s protocol. Paired-end Fastq reads were trimmed for quality and mapped to the published sequence for the SARS-CoV-2 reference isolate Wuhan-Hu-1 (RefSeq: NC_045512.2) using CLC Genomics Workbench version 21 from which consensus sequences were generated. Stocks were confirmed to be free from contamination by adventitious agents by analysis of reads that did not map to SARS-CoV-2 or cell-derived sequences.

Virus samples were titrated using a 50% Tissue Culture Infectious Dose (TCID_50_) assay. In brief, samples were serially 10-fold diluted in DMEM supplemented with 2% v/v FBS, 2 mM GlutaMAX supplement, 100 U/mL penicillin, and 100 μg/mL streptomycin, starting at a 1:10 dilution. For each dilution in the series, six replicate wells were prepared per sample in 96-well plates (50 µL per well) into which 2 × 10^4^ Vero E6 cells/well in 100 µL volume were added. Plates were incubated at 37 °C/5% CO_2_ for four days before being assessed for the presence of cytopathic effect. TCID_50_ titres were calculated using the six replicates for each sample and the Spearman-Kärber method [21].

### Preliminary Infection Studies with Human Nasal Epithelial Cells

Prior to infection of the HNE cells, the apical face of each transwell culture was washed twice with 300 μl Dulbecco’s Phosphate Buffered Saline containing Calcium and Magnesium ions (DPBS+; Thermo Fisher Scientific) to remove accumulated mucus. Basal media containing either a 1:1,000 dilution of remdesivir stock (5 mM final concentration) or the equivalent volume of DMSO vehicle was prepared in PneumaCult-ALI medium (STEMCELL Technologies) and was used to replace media in the appropriate wells of the 24-well plates containing the transwells. Mock and virus-only treated cells received the media with DMSO, while the drug-only and virus+drug cells received the media with 5 μM remdesivir. The transwells were incubated at room temperature for one hour before being infected with an apical administration of 100 μl SARS-CoV-2 Delta VOC inoculum at an MOI of 0.01, or 100 μl media without virus, for a further hour at 37°C/5% CO_2_. After the 1hr incubation, 200 μl DPBS+ was added to the apical side of each well before being removed along with any inoculum. The plates were returned to incubate at 37°C/5% CO_2_, and a back titration was performed to confirm the inoculum titre.

Basal media and apical wash samples were harvested daily from quadruplicate wells on days 1-4 post-infection. 350 μl DPBS+ was added to the apical side of the appropriate wells, and the plates were incubated at 37°C/5% CO_2_ for 30 min to allow for equilibration of the samples. After incubation, the apical washes were removed and stored at -80°C until titration by TCID_50_ assay. 1 ml samples of basal media were also removed from each well and stored at -80°C until titration.

Wells for harvest on Day 3 and 4 post-infection received a media/treatment change on Day 2 post-infection. DMSO- and remdesivir-containing media was prepared as described above and was used to replace the media in the 24-well plates, as appropriate.

### Preliminary Infection Studies with Human Cardiomyocytes

Previously seeded 12-well plates of attached, differentiated cardiomyocytes were treated with 1 ml STEMdiff Cardiomyocyte Maintenance Medium (STEMCELL Technologies) containing 1:1,000 diluted remdesivir (5 mM final concentration) or 1:1,000 dilution of DMSO vehicle. The plates were incubated at room temperature for an hour before the media was removed and the cells were treated with 200 μl media containing the remdesivir or DMSO, along with SARS-CoV-2 Delta VOC inoculum at an MOI of 0.01 for infected wells, for a further hour at 37°C/5% CO_2_. After the 1h infection, 400 μl DPBS+ was added to each well before being removed along with any inoculum. 1ml medium containing remdesivir or DMSO was added to each well as appropriate and plates were returned to incubate at 37°C/5% CO_2_. A back titration was performed to confirm the inoculum titre.

1ml aliquots of media harvested daily from quadruplicate wells on days 1-4 post-infection and stored at -80°C until titration by TCID_50_. Media in the Day 3 and 4 wells was replaced on Day 2 post-infection with media containing the appropriate additive (remdesivir or DMSO).

### Drug Selection, Procurement, and Preparation

Prospective drugs were down-selected from the Compounds Australia Open Drug collection using a set of filters described previously [12]. Based on the results of our previous study using a human airway tissue model, fluvoxamine, everolimus, pyrimethamine, aprepitant, and sirolimus were tested alongside the control drugs remdesivir, molnupiravir, and nirmatrelvir (PF-07321332; the active ingredient in Paxlovid) [13]. All of the drugs were obtained from Selleck Chemicals (Houston, TX, USA) or Sigma Aldrich (St Louis, MO, USA). Where possible, drugs were obtained pre-dissolved as 10 mM stocks in DMSO. For drugs not available in this format, 10 mM DMSO stocks were prepared and sterilised by filtration through a 0.22 µm syringe filter. As ondansetron was insoluble in DMSO, it was dissolved in 10 mM HCl and then filter sterilised.

### Drug Antiviral Efficacy Testing with Human Cardiomyocytes

Dilution ranges of the test and control drugs were prepared in deep well plates in STEMdiff Cardiomyocyte Maintenance Medium (STEMCELL Technologies). Fluvoxamine, everolimus, pyrimethamine, aprepitant, sirolimus, and the control drug molnupiravir were prepared to final concentrations of 25, 10, 4, 1, and 0.4 μM, while the other two control drugs, remdesivir and nirmatrelvir, were prepared to final concentrations of 10, 4, 1, 0.4, and 0.05 μM. These included 2x concentration preparations for infection wells so that the correct final concentrations could be obtained when combined 1:1 with prepared inoculum. Media for negative control and virus-only wells contained the equivalent amount of DMSO as the 25 μM drug concentrations.

The media was removed from differentiated cardiomyocytes, previously seeded into 12-well plates, and replaced with 0.5ml of the appropriate treatment medium. The plates were incubated at room temperature for an hour before the media was removed and the cells were treated with 200 μl media containing the appropriate treatment, along with SARS-CoV-2 Delta or Omicron VOC inoculum at an MOI of 0.01 for infected wells, for a further hour at 37°C/5% CO_2_. After the 1h infection, 400 μl DPBS+ was added to each well before being removed along with any inoculum. 1ml medium containing the appropriate treatment was added to each well and plates were returned to incubate at 37°C/5% CO_2_. A back titration was performed to confirm the inoculum titre. 1ml aliquots of media were harvested on Day 2 post-infection and stored at -80°C until titration by TCID_50_.

### Immunofluorescence and Confocal Microscopy

For the preliminary infections, HNE cells in transwells and cardiomyocytes cultured on treated coverslips representing each study condition were fixed and processed for immunofluorescence confocal microscopy as previously described [22]. Briefly, cells were washed three times with DPBS+ at room temperature, and fixed with 4% w/v paraformaldehyde (Electron Microscopy Sciences, Hatfield, PA, USA) for a minimum of 60min at room temperature. The fixative was aspirated and neutralised with 100mM glycine in DPBS+ for 10min at room temperature. Cells were incubated with permeabilisation buffer (PB; 0.5% v/v Triton-X100 in DPBS+) for 30min on ice. For HNE transwells, the PB was washed off, and the filters were excised from the inserts using a sharp scalpel, cut in half (for test and control primary antibodies), transferred to Eppendorf tubes, and incubated for 90min at 4°C in immunofluorescence buffer (IB; DPBS+ containing 0.1% w/v bovine serum albumin, 0.2% v/v Triton-X100, and 0.05% v/v Tween-20) containing 10% v/v normal goat serum (block buffer; BB). For the cardiomyocytes, the coverslips were placed in fresh 24-well plate wells before having the PB washed off, and BB added for 90min incubation at 4°C. For both cell types, at the end of the 90min incubation, the BB was removed and replaced with primary antibody (antibody information can be found in Table S1) diluted in BB before being returned to 4°C for 48h. Following the incubation, the primary antibody was washed off three times with IF buffer, 5min per wash, at room temperature. Flurophore-conjugated secondary antibody and Hoechst stain, diluted in PB, was added to each sample, and incubated for 3h at room temperature. Secondary antibody was washed off five times with IF buffer, 5min per wash. Filters were transferred to slides, incubated at room temperature for 30min with DAPI, and washed once with PBS before being mounted in FluoroSave reagent (Merck Millipore, Burlington, MA, USA). Coverslips were treated similarly before being sealed with nail polish. Confocal microscopy imaging was acquired using a Zeiss LSM 780 system. The acquired Z-sections were stacked and processed using ImageJ software (National Institutes of Health, Bethesda, MA, USA). Orthogonal views were generated using ZEN 3.1 software from Zeiss Microscopy (Oberkochen, Germany).

## Results

### Preliminary Infections of Tissue Models

To evaluate additional tissue models for their suitability for use in the testing of antiviral efficacy of drugs against SARS-CoV-2 infection, we assessed an ALI HNE model and a stem cell-derived cardiomyocyte model. Nasal inferior turbinate brushings were collected, conditionally reprogrammed in cell expansion culture, and used to generate HNE cultures as previously described [19]. The resulting transwell cultures consisted of a multi-layered pseudostratified columnar epithelium with a uniform cobblestone morphology. Beating cilia were also observed, as is characteristic of mucociliary differentiation (data not shown), and well-developed apical cilia were detected by staining for acetylated α-tubulin (**Figure 1a**). In addition, differentiated cardiomyocytes were generated from embryonic stem cells, which showed the characteristic collective beating phenotype by Day 11 of differentiation (**Supplementary Video 1**). Cardiomyocyte differentiation was further confirmed by immunostaining for sarcomeric α-actinin and cardiac troponin T in Day 20 cultures compared to undifferentiated stem cells, with most cells positive for both markers (**Figure 1b**).

**Figure 1:**
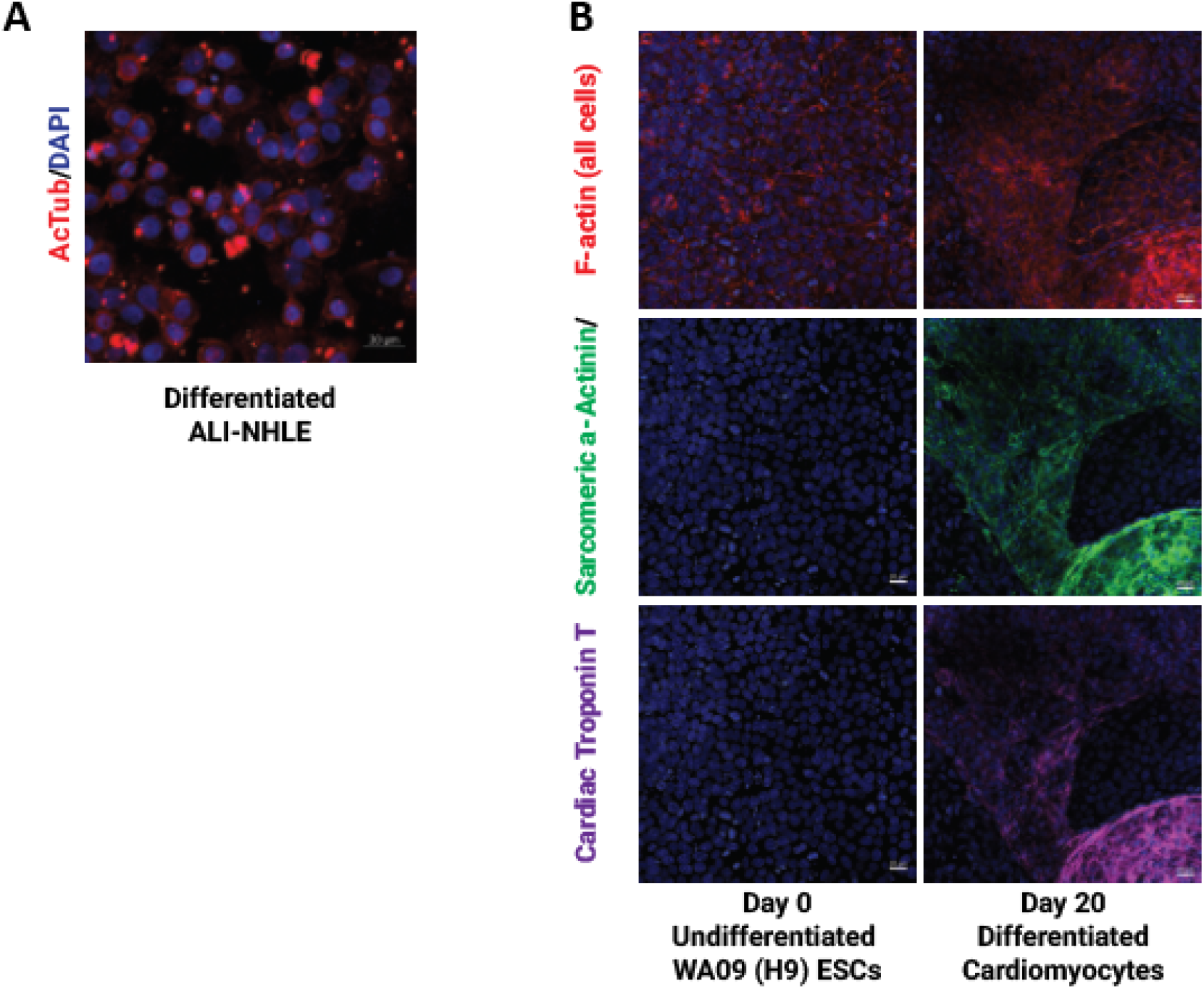
Differentiation of Human Nasal Epithelium (HNE) and Cardiomyocyte Models. A) Immunofluorescent staining of acetylated tubulin (red) to detect apical cilia with DAPI (blue) staining of cell nuclei. B) Immunofluorescent staining of undifferentiated WA09 embryonic stem cells or Day 20-differentiated cardiomyocytes stained for f-actin (phalloidin; red), DAPI (blue), and cardiomyocyte markers sarcomeric α-actinin (green) and cardiac troponin T (purple). Scale bar represents 20µm.

In order to determine the suitability of the HNE and cardiomyocyte tissue models for use with SARS-CoV-2, preliminary infection studies were performed to assess the growth of the SARS-CoV-2 Delta VOC in the presence and absence of 5 μM remdesivir. This concentration of remdesivir had been previously selected for use in airway and alveolar tissue models based on a review of the literature and was used in this study for consistency [13]. Sufficient HNE and cardiomyocyte culture wells were prepared for four treatment conditions (mock, drug-only, virus-only, and virus+drug) to allow quadruplicate sample sets to be harvested for each condition on Days 1-4 post-infection. Viral infection was observed by fixed-cell immunofluorescence in HNE cultures (**Supplementary Figure 1**), and by the generation of cytopathic effect in cardiomyocytes (infected cardiomyocytes detach from treated glass coverslips).

Samples collected from the cardiomyocyte culture medium, and HNE basal media and apical washes at each timepoint were titrated to determine the viral loads. For the cardiomyocytes, titres in the virus-only wells increased rapidly, peaking on Day 2 with values between 10^5^ and 10^6^ TCID50/ml and slowly declining thereafter (**Figure 2a**). For the HNE cells, viral growth in the virus-only wells was also rapid, reaching titres of 10^4^-10^6^ TCID_50_/ml by Day 2 (albeit with substantially higher variation than with the cardiomyocytes) and maintaining those titres on Days 3 and 4 (**Figure 2b**). Infectious virus was only detected in apical wash samples, with all basal medium samples below the limit of detection (10^2^ TCID_50_/ml).

**Figure 2:**
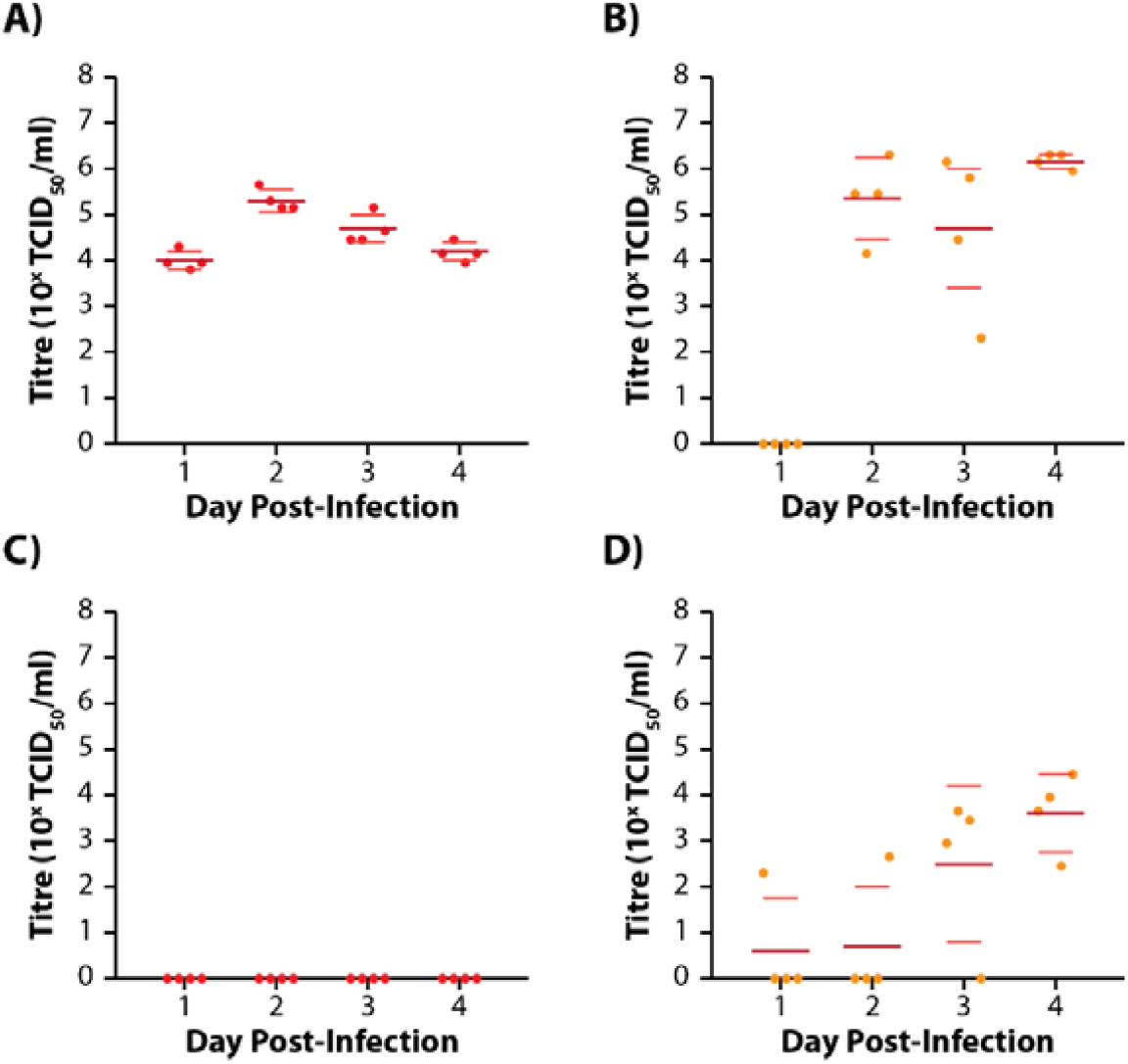
SARS-CoV-2 Infection and Response to 5µM Remdesivir in HNE and Cardiomyocyte Models. Viral titres of SARS-CoV-2 Delta variant in media and apical washes of (A) cardiomyocyte and (B) HNE cultures, respectively, without drug treatment. 5µM remdesivir was effective at suppressing detectable infection in (C) cardiomyocyte cultures, as well as retarding growth in (D) HNE cultures until Day 3 post-infection. Infectious virus was not detected in basal media samples from HNE culture. Dots represent virus titres from individual quadruplicate samples. Horizontal bold lines represent the mean titre of replicates, while lighter coloured lines represent the standard error of the mean (SEM).

The addition of 5 μM remdesivir to the culture media prevented viral growth in cardiomyocytes (**Figure 2c**), but appeared to only impede growth in HNE cultures. Indeed, by Day 4 post-infection, all four HNE culture replicates had detectable virus (titres 10^2^-10^4^ TCID_50_/ml) in apical wash samples in the presence of 5 μM remdesivir (**Figure 2d**). Next-generation sequencing of the Day 4 HNE samples revealed that the populations of virus in these samples had no consensus-level mutations compared to the input stock virus indicating that this was not an adaptation of the virus to the drug.

### Testing of Drug Anti-SARS-CoV-2 Efficacy in Human Cardiomyocyte Culture

Given the significant variability in virus titres observed with the HNE cultures, the decision was made to use the cardiomyocyte model for drug antiviral efficacy testing. Unfortunately, due to constraints on the number of wells available and the complexities with handling cultures under BSL-4 conditions (required at the time for work involving infectious SARS-CoV-2 at our facility), a trade-off had to be made between having multiple concentrations of each drug and having replicate samples. A decision was therefore made to focus on the range of concentrations in this screening assay to minimise the risk of discounting potentially active compounds.

Five test drugs - fluvoxamine, everolimus, pyrimethamine, aprepitant and sirolimus - and three control drugs - remdesivir, molnupiravir, and nirmatrelvir (PF-07321332) - were tested at five concentrations (25, 10, 4, 1, 0.4 μM for test drugs and molnupiravir; 10, 4, 1, 0.4, 0.08 μM for remdesivir and nirmatrelvir) against both Delta and Omicron VOC. Triplicate mock and virus-only wells were also included as controls. Media samples were harvested 2 days post-infection (corresponding with peak viral titre in the preliminary infection assay) and were titrated to determine virus loads.

Both Delta and Omicron VOC grew well in the cardiomyocytes, with Delta titres around 10^4^ TCID_50_/ml and Omicron titres around 10^5^ TCID_50_/ml in the virus-only wells. For both variants, all concentrations of remdesivir tested completely inhibited viral growth (**Figure 3a**). Nirmatrelvir reduced viral titres around 100-fold at 0.08 μM and inhibited growth entirely at higher concentrations (**Figure 3b**). Although having a similar mechanism of action to remdesivir, molnupiravir failed to inhibit both Delta and Omicron growth at 0.4 and 1 μM concentrations, however was effective at preventing Delta growth at concentrations of 4 μM and higher (**Figure 3c**). Interestingly, Omicron was able to grow in the presence of 4 μM monupiravir although to a titre 100-fold lower than the virus-only control wells.

**Figure 3:**
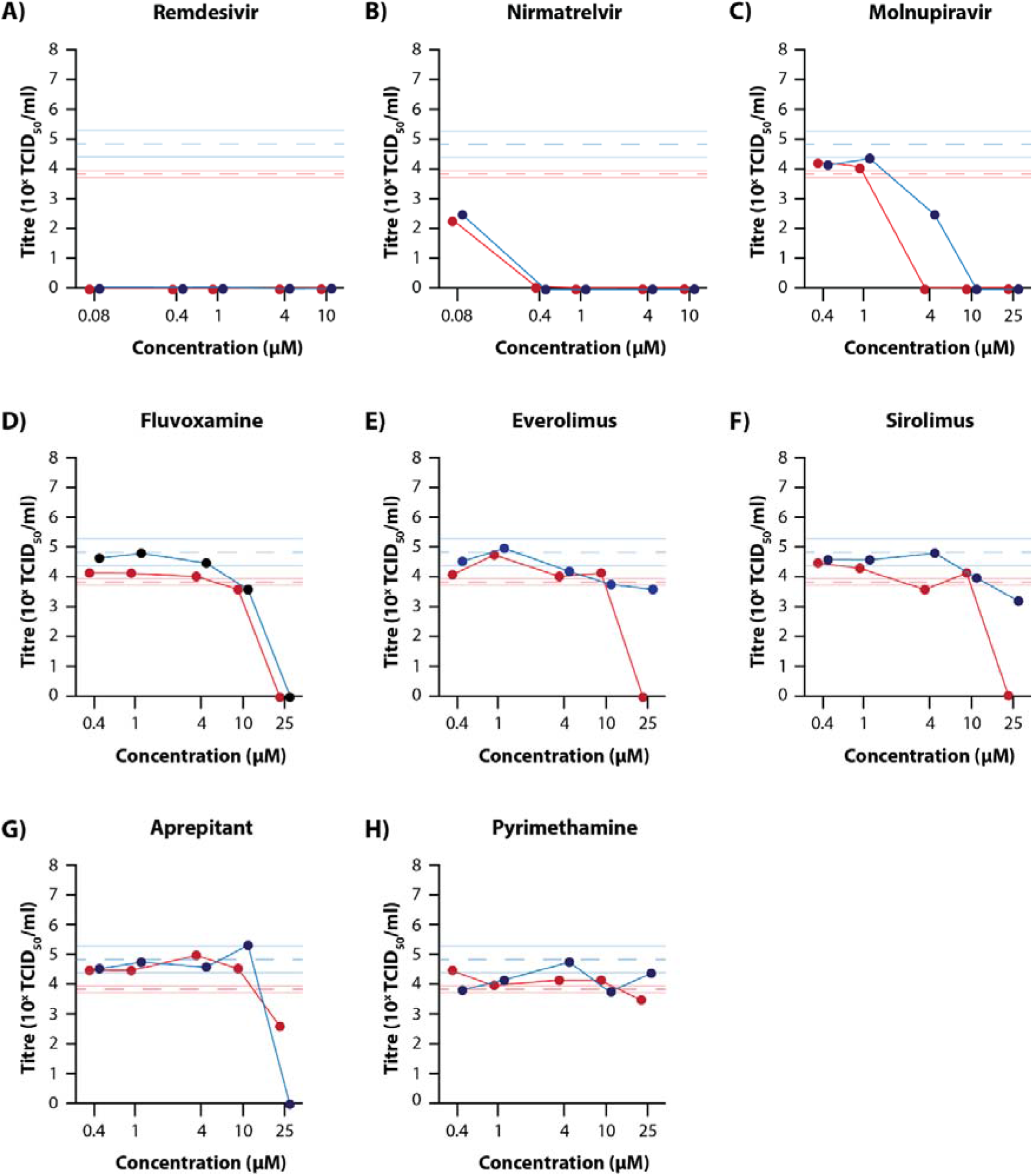
Antiviral Efficacy of Selected Drugs Against SARS-CoV-2 Delta and Omicron in the Cardiomyocyte Model. Differentiated cardiomyocytes were infected with a 0.2 MOI of Delta (red) or Omicron (blue) SARS-CoV-2 variant in the presence of 0.08, 0.4, 1, 4, 10 µM remdesivir (A) or nirmatrelvir (B), or 0.4, 1, 4, 10, 25 µM molnupiravir (C), fluvoxamine (D), everolimus (E), sirolimus (F), aprepitant (G), or pyrimethamine (H) with media samples collected and titrated after 48h. The red and blue dashed lines represent mean titre from three virus-only wells for Delta and Omicron, respectively. Pale red and blue lines either side represent SEM for the virus-only titres.

Against both Delta and Omicron VOC, fluvoxamine prevented virus growth at the highest concentration (25 μM), but not at 10 μM or lower (**Figure 3d**). A similar complete inhibition was observed for Delta with 25 μM everolimus and sirolimus, but interestingly this inhibition did not extend to Omicron (**Figure 3e & 3f**). Conversely, 25 μM aprepitant completely inhibited Omicron growth, while only reducing Delta titres <100-fold (**Figure 3g**). Pyrimethamine failed to reduce viral titres of either variant at any concentration tested (**Figure 3h**).

## Discussion

The ability to screen approved drugs for efficacy against off-target diseases is important in a fast-paced pandemic situation, as it can potentially identify effective therapies that have already undergone rigorous safety testing and get them to the clinic faster than novel treatments. However, given the thousands of drugs that have been approved by regulatory bodies around the world, screening for efficacy is difficult. Several studies have aimed to use high-throughput tissue culture-based approaches for screening of approved drugs against SARS-CoV-2 infection (e.g. [2, 3, 6]), however the behaviour of both the virus and drugs can differ significantly in tissue culture compared to human tissues, raising the question of how helpful such screening studies are. Indeed, as seen with chloroquine/hydroxychloroquine and ivermectin, the ability to prevent viral-induced cell death in tissue culture does not mean that the drugs will be effective in treating the disease in patients [4, 5, 9, 10]. Moreover, the exploitation of such *in vitro* studies can result in the misuse of ineffective drugs with potentially lethal results.

Although they have some drawbacks with regards to cost and ease of handling, 3D tissue culture models represent a more suitable approach to drug repurposing studies, as they better recapitulate the cell types and structures found within host tissues [23]. Furthermore, their suitability can be further improved by selecting models that represent key tissues targeted by the virus. For viruses like SARS-CoV-2, that appear to have a broad tissue tropism, the use of a panel of multiple tissue models is important to ensure that any antiviral efficacy is broad and not limited to particular tissue types.

Cardiomyocytes and cells of the nasal epithelium have both been demonstrated to be important sites of replication during COVID-19, contributing to virus transmission and virus-induced cardiac disease [14, 18]. Accordingly, we selected these tissue models to increase the repertoire of possible models for drug repurposing studies. As expected, both tissue models were permissive to SARS-CoV-2 infection, with high viral titres attained by Day 2. Whereas viral titres in the cardiomyocytes started to decrease on Days 3 and 4 post-infection, high titres persisted in the nasal epithelial models from Day 2 to 4, results that are in agreement with previous studies that have shown nasal epithelial cells to sustain persistent SARS-CoV-2 infection despite prolonged antiviral responses [17]. Furthermore, while 5 μM remdesivir was sufficient to prevent SARS-CoV-2 infection in cardiomyocytes (as well as airway and alveolar tissue models, shown in our previous study [13]), the same antiviral treatment of nasal epithelial cells failed to prevent SARS-CoV-2 infection, with all four replicate wells positive for infectious virus by Day 4 post-infection. Given that sequencing of these drug-escape isolates revealed no consensus-level sequence changes relative to the input virus stock, this appears to potentially be a feature of the nasal epithelium being able to support viral replication in the presence of antiviral factors. This is supported by other studies showing that nasal epithelial cells support persistent infection in the presence of antiviral immune responses (human tissue model; [17]) and neutralising antibodies (mouse vaccine challenge model; [18]).

In our previous study, we used a commercially available *ex vivo* human airway model to assess approved drugs for anti-SARS-CoV-2 activity following an *in silico* down-selection process from an original list of nearly 8,000 compounds [12, 13]. This study demonstrated that, against Omicron, the selective serotonin reuptake inhibitor fluvoxamine demonstrated antiviral efficacy at 25 μM, while against Delta the results at 25 μM varied between experiments. The other drugs tested (everolimus, pyrimethamine, aprepitant, and sirolimus) showed minimal efficacy at the concentrations used against both Delta and Omicron VOC. In this study, the same drugs were tested with the cardiomyocyte tissue model with fluvoxamine preventing both Delta and Omicron growth at 25 μM. Unlike the airway cells, everolimus and aprepitant also fully inhibited Delta (but not Omicron) growth at a 25 μM concentration, while 25 μM aprepitant prevented Omicron growth and reduced Delta titres by <100-fold. Responses in both the airway and cardiomyocyte models against both Delta and Omicron were very similar for nirmatrelvir (PF-07321332; the active ingredient in Paxlovid) and molnupiravir. Interestingly, the growth of both variants appeared to be more affected by remdsesivir in the cardiomyocyte model than in the airway model with complete inhibition of viral growth with 0.08 μM drug in cardiomyocytes compared to 4 μM in the airway model – a 20-fold difference in concentration.

Of the drugs tested, only fluvoxamine appeared to have antiviral efficacy against both the Delta and Omicron variants, however such efficacy was only observed at 25 μM. Such a high concentration is not achievable in patients, where plasma concentrations are usually around 0.3 μM during treatment [24]. This may explain why, although apparently capable of inducing an antiviral effect, clinical trials of fluvoxamine for the treatment of COVID-19 have failed to show clear antiviral efficacy [24-26].

Although this study did not identify an effective, repurposed drug for the treatment of COVID-19, it demonstrates that (a) multiple tissue models can be successfully used for the assessment of repurposed drugs against SARS-CoV-2 infection; and our methodology could be useful for (b) future coronavirus outbreaks, as well as (c) screening compounds in libraries such as ChEMBL (for example [27]). Comparison of remdesivir efficacy against SARS-CoV-2 infection in cardiomyocytes with efficacy in the airway model used previously ([13]), clearly demonstrates that the effective concentrations of drugs can differ significantly in different tissue types highlighting the need for panels of differing tissue models to be used for thorough efficacy testing. Studies like ours help to advance the implementation of the FDA Modernization Act which “authorizes the use of certain alternatives to animal testing, including cell-based assays and computer models, to obtain an exemption from the Food and Drug Administration to investigate the safety and effectiveness of a drug”, and “removes a requirement to use animal studies as part of the process to obtain a license for a biologic” [28].

## Funding

This research was kindly funded (Principal Investigator: S.S.V.) by the Australian Department of Health through its Medical Research Future Fund (Grant Number MRF2009092) and the United States Food and Drug Administration (FDA) Medical Countermeasures Initiative (Contract Number 75F40121C00144). Establishment of the ALI-HNE was funded by philanthropic funding from the Kim Wright Foundation awarded to E.V. The salaries for E.V. and B.M.T. was supported by an NHMRC Ideas grant awarded to EV and BMT (APP1181580). The article reflects the views of the authors and does not represent the views or policies of the funding agencies, including the FDA.

### Institutional Review Board Statement

This study involved no human or animal subjects; however, per the regulations in Australia, the use of human embryonic stem cells (for cardiomyocyte production) and the infection of both sets of tissue models were reviewed and approved by the CSIRO Human Research Ethics Committee (Approval Number: 2021_083_LR; Approval Date: 14 September 2021). This was in addition to study approval for the generation of human nasal epithelial cells from the Sydney Children’s Hospital Network Ethics Review Board (HREC/16/SCHN/120) and the Medicine and Dentistry Human Ethics Sub-Committee, University of Melbourne (HREC/2057111).

### Informed Consent Statement

Written consent was obtained from all participants prior to collection of biospecimens from which the de-identified, human nasal epithelial cells were derived. Informed consent is not applicable for the WA09 embryonic stem cells which were obtained from WiCell, Madison, WI, USA.

## Data Availability Statement

Any underlying data not presented can be provided by the corresponding authors upon reasonable request.

## Acknowledgements

The authors would like to acknowledge the contributions made by the broader ‘sySTEMs initiative’ project members.

